# Early-Stage Cultures of *Acidithiobacillus ferrooxidans* Enable Efficient Bioleaching of Li-ion Battery Cathode Material

**DOI:** 10.64898/2026.08.04.742857

**Authors:** Brooke E. Elander, Mengyun Jiang, Cameron Guthrie, Zachary Ibrahim, Babak Momeni, Dunwei Wang

## Abstract

In response to the growing need to recycle E-waste, specifically that of lithium-ion batteries (LIBs), the development of sustainable recycling methodologies will be vital. As one of the most sustainable and low-cost options, biohydrometallurgy (BioHM) uses biological organisms to facilitate the recovery of critical metals from spent LIBs. Despite the advantages of BioHM, its slow kinetics, due to the reliance on the metabolic activity of microorganisms, limit its large-scale application for the closed-loop recycling of LIBs. In this work, we investigate the correlation between the incubation time of Acidithiobacillus ferrooxidans (Atf) and its leaching efficiency of four common elements in LIBs: Li, Ni, Mn, and Co. We assess how specific incubation times along the biological growth curve affect the leaching efficiency of Li[Ni0.6Mn0.2Co0.2]O2 (NMC622), a model electrode material. Our results show that pH alone is not an accurate descriptor of the leaching efficiency of Atf cultures used at different growth stages. The addition of NMC622 during the bacterial lag phase reaches similar or even improved extraction compared to cultures used after reaching the exponential or stationary phases. A simple model of leaching dynamics shows predictions consistent with our experimental observations under the condition that the inhibition of bacterial growth by NMC is not severe. Our findings indicate that early-stage cultures can alleviate the kinetic bottleneck and improve the throughput of recovering critical materials from spent batteries.

## 1. Introduction

The global energy demand has propelled the utilization of fossil fuels at an unsustainable pace.^1,2^ In an effort to mitigate the detrimental effects from carbon dioxide emissions, the widespread implementation of lithium-ion batteries (LIB) represents a transition toward renewable energy and energy storage systems.^3,4^ In 2023, total global battery usage surpassed 2,400 gigawatt-hours (GWh), with LIBs powering over 40 million electric vehicles.^5^ With such growth, however, there has been a resulting accumulation of waste from spent batteries, which may pose devastating impact on the environment when left untreated.^6–8^ To address this potential challenge and to recover critical minerals such as lithium (Li), cobalt (Co), and nickel (Ni), researchers have turned to recycling spent LIBs as a critical strategy.^9^ Compared to primary metal extraction, recycling provides significant environmental benefits by lowering greenhouse gas emissions, energy use, and water consumption.^10^ Despite these benefits, modern industrial recycling methods rely on hazardous chemicals and costly infrastructure, which makes it exceedingly challenging to scale up these approaches in a sustainable manner.^11,12^ In recognition of this limitation, recent research has shifted to recycling strategies alternative to traditional pyrometallurgy (PM)^13^ and hydrometallurgy (HM)^14^, aiming to decrease reagent and energy use while enhancing metal selectivity in an cost-effective manner.

While HM is highly effective in metal recovery, it requires high volumes of concentrated acids, bases, or potentially hazardous reagents such as hydrogen peroxide (H_2_O_2_).^15^ Biohydrometallurgy (BioHM) shares core principles with HM but utilizes the functionality of living organisms to recover metals from raw materials or various waste streams.^15^ In BioHM, living organisms, typically bacteria or fungi, drive leaching through the generation of acids, oxidants, or metal binding compounds that mobilize metals into solutions from which they can be separated and purified. Bacterial species such as *Acidithiobacillus ferrooxidans* (*Atf*) and *Leptospirillum ferrooxidans* are the classical bioleaching workhorses for sulfide ores.^16^ Heterotrophic and acid tolerant fungi such as *Aspergillus niger*, *Penicillium spp*., and other filamentous fungi have also been considered for rare earth elements (REEs) and technology wastes.^17^ Possible advantages of BioHM include its reduced chemical demand, emissions, and energy input, making BioHM a potentially safer and more cost-effective alternative to traditional HM approaches.^18^ BioHM has already been applied industrially to recover copper^19^ and gold^20^ from ore and is now being explored to extract critical and REEs^21^ from both natural deposits and secondary waste such as electronics and electric vehicle (EV) components. Because microbes grow and act at ambient temperature and pressure, BioHM can be applied to low grade ore or fine heterogeneous waste streams where conventional pyrometallurgy (PM) is inefficient or costly.

*Acidithiobacillus ferrooxidans* (*Atf*), a chemolithotrophic bacteria, is commonly chosen for BioHM. Over its growth period, *Atf* oxidizes elemental sulfur and ferrous iron (Fe^2+^), producing sulfuric acid (H_2_SO_4_) and ferric iron (Fe^3+^) as byproducts. The biogenically produced H_2_SO_4_ dissolves various metal oxides and sulfides, enabling the recovery of critical metals in spent LIBs such as Li and Co. Within this system, Fe^2+^ functions as a reducing agent and Fe^3+^ as an oxidizing reagent, together driving redox reactions which convert metals into soluble forms.^22^ This process enables metal recovery without the need for expensive and hazardous reagents associated with HM^23^, creating an environment-friendly process.^24^ However, an important challenge for BioHM is the inherently slow reaction kinetics of both cellular growth and bioleaching.^25^ Microbial metabolic activity results in slower metal solubilization rates compared to other leaching methods, and these rates are influenced by factors such as pH, redox potential, metal toxicity, and temperature.^26^ In industrial applications, reduced substrate availability and reduced oxygen availability in bioreactors may also limit reaction rates, by mechanisms like jarosite formation on substrate surfaces.^27^ The efficiency of REE bioleaching, for example, is limited by the slow kinetics of REE extraction by organisms.^28^ In a fungal bioleaching experiment, the reaction time averaged between 15 and 60 days, which is considerably longer than that of conventional HM and PM operations and underscores the inherently slower kinetics of the bioleaching process.^25^ Additionally, few studies have explored the kinetics of bioleaching reactions; this has made it hard to compare leaching performances across different investigations.^29^ Collectively, these observations highlight the kinetic bottleneck and the need for more systematic kinetic studies to optimize processes and allow reliable comparison of results across different BioHM systems.

In this study, we systematically investigate the leaching efficiency of Li[Ni_0.6_Mn_0.2_Co_0.2_]O_2_ (NMC_622_) electrode powder across different incubation periods of the *Atf* culture. Our data reveals how leaching efficiencies are affected by the growth stage of the bacterial cultures. We show that early addition of NMC material when cultures are in the lag phase leads to leaching efficiencies comparable to those of exponential or stationary phases. Our findings suggest that exposure to NMC early in the growth process does not inhibit the growth and metabolic activities of *Atf*, allowing efficient leaching in an overall shorter timeframe.

## 2. Materials and Methods

### 2.1. Microorganism and Growth

The *Acidithiobacillus ferrooxidans* strain F221 (DSM1927) was obtained from DSMZ, Germany. The bacteria were cultivated in a M9K medium^30^, with components (NH_4_)_2_SO_4_ (3.0 g/L) (Millipore Sigma), KCl (0.1 g/L) (Fisher Scientific), K_2_HPO_4_·3H_2_O (0.5 g/L) (Fisher Scientific), MgSO_4_ ·7H_2_O (0.5 g/L) (Fisher Scientific), Ca(NO_3_)_2_ (0.01 g/L) (Fisher Scientific), and FeSO_4_·7H_2_O (44.22 g/L) (STREM). The *A. ferrooxidans* culture was used as the parent culture to inoculate subsequent samples once they reached the stationary phase.

**Table 1.** The components of the M9K media used for the culture system.

| MEDIA COMPONENT | COMPOUND | QYT. (g/L) |
| --- | --- | --- |
| Nitrogen source | $(\text{NH}_4)_2\text{SO}_4$ | **3.0** |
| Magnesium source | $\text{MgSO}_4 \cdot 7\text{H}_2\text{O}$ | **0.5** |
| Potassium source | KCl | **0.1** |
| Phosphate source and buffer | $\text{K}_2\text{HPO}_4 \cdot 3\text{H}_2\text{O}$ | **0.5** |
| Calcium source | $\text{Ca}(\text{NO}_3)_2 \cdot 4\text{H}_2\text{O}$ | **0.01** |
| Energy source | $\text{FeSO}_4 \cdot 7\text{H}_2\text{O}$ | **44.2** |

The 24-well experiments were conducted in a 24-well flat-bottom microtiter plate (Falcon™ 353047) setting with 1.5 mL of M9K media in each sample. All samples’ pH’s were adjusted to 1.6-1.8 using 2M H_2_SO_4_ (Millipore Sigma) for the initial acidification step. Each sample was inoculated with 10% v/v parent culture and went through incubation at 30 °C under shaking at 170 rpm. The 24 wells were divided into 6 groups: early lag phase (**EL, 2h incubation**), late lag phase (**LL, 4h incubation**), early exponential phase (**EE, 8h incubation**), mid exponential phase (**ME, 12h incubation**), late exponential phase (**LE, 24h incubation**), and early stationary phase (ES, 48h incubation), with 4 replicates each. Biological studies, including [Fe^3+^], oxidation reduction potential (ORP), and pH measurements, were performed pre- and post-addition of cathode materials. When each group of 4 wells reached their set incubation time, NMC_622_ was added (pulp density of 10 g/L) and allowed to undergo leaching for another 48 hours. Biological studies were taken every 24 hours after cathode material was added. For example, this would mean 2h samples from inoculation to analysis of metal concentration would take 50 hours, and 4h samples would take 52h. In addition, control experiments were carried out in the same culture media and setting without bacteria.

The test tube experiments were conducted in glass test tubes (Fisherbrand™ Disposable Borosilicate Glass Tubes - O.D. x L: 12 x 75mm, #14-961-26) setting with 5 mL of M9K media in each sample. All other parameters and conditions were identical to the 24-well setting.

### 2.2. Bioleaching Studies

Given the allotted incubation periods, the bioleaching experiments were conducted by analyzing [Fe^3+^], pH, ORP, and biogenic acid at each point before the addition of cathode materials (pulp density: 10 g/L). The same studies were then conducted at 24h increments for the rest of the 48h leaching timeframe. This resulted in a two-step process (allowing for different bacterial growth phases to be reached before adding cathode material) that took place in one pot. For all leaching experiments, the processes were conducted at 30°C with continuous shaking at 170 rpm. After 48h of leaching, the resulting liquor was filtered from residual materials in the solution using a PVDF membrane filter with polypropylene housing and a nominal pore size of 0.22 *μ*m. The quantities of the critical metals (Li, Ni, Mn, and Co) in the filtered leaching liquor were analyzed using inductively coupled plasma optical emission spectroscopy (ICP-OES). The leaching efficiency of the metals was determined by comparing the samples with an *aqua regia* digestion (HCl:HNO_3_ = 3:1) of the pristine cathode materials (**Equation 1**), which was achieved by adding the same pulp density (10 g/L) ratio of cathode material (NMC_622_) to an *aqua regia* solution and autoclaving at 90°C for 12 h.

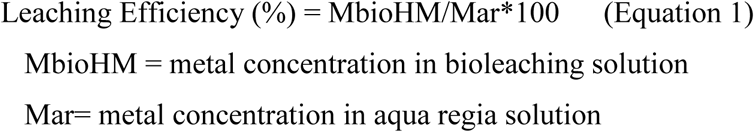

### 2.3. Analytical Methods

#### 2.3.1. Quantification of ferrous iron concentration

[Fe^3+^] was quantified by measuring the optical absorption using a microplate reader (BioTek Epoch 2). A 10% 5-sulfosalicylic acid (SSA) (Lab ally) solution was used as an indicator. 2 μL of a given culture was collected and diluted with 968 μL of DI water, followed by mixing with 30 μL of SSA. The spectrophotometric optical absorption was measured at a wavelength of λ= 500 nm.

#### 2.3.2. Analytical instruments

ORP and pH were measured using an Orion Star A211 Benchtop pH Meter (Thermo Scientific, STARA2110). For ORP, a Thermo Scientific™ Orion™ Redox/ORP/Temp Electrode (#9179BNMD) was used at 25°C. To measure the pH, we used a Thermo Scientific™ Orion™ Double Junction Semi-Micro Stem Glass Bodied Combination pH Electrode (#9110DJWP). The concentration of the metals of interest was taken using an ICP-OES (Thermo scientific)

#### 2.3.3. Biogenic acid

The concentration of the produced biogenic acid of H_2_SO_4_ was taken using the classical acid-based titration method. In a typical experiment, a sample of the culture was diluted 10-fold and removed of any cells and debris though centrifugation at 12,000 rpm for 10 min. Using a 0.1 M NaOH solution and a bromothymol blue indicator the solution was titrated, and acid concentration was quantified.

### 2.4. Mathematical Modeling of Bioleaching Dynamics

All simulations were implemented in Matlab R2025b. Dynamics were simulated using a simple forward Euler recipe, with a time step of 0.001 h. To model different cases, all growth simulations started from the same initial density of the bacterial cells, but NMC622 was added to the bacterial cultures at different time points of EL, LL, EE, ME, LE, and ES (matching the experimental protocol). The duration of simulated dynamics was 96 hours.

## 3. Results and Discussion

A barrier to the implementation of biohydrometallurgy processes for LIB recycling has been the slow kinetics of the organisms involved. While the predominant metabolic pathway of *Atf* is the oxidation of ferrous iron to ferric (Fe^2+^ → Fe^3+^ + e^-^), which is consequently used as a proxy to indicate cell activity and growth, the overall process involves additional metabolic reactions that influence metal solubilization. One such process is the biogenic formation of H_2_SO_4_. The generation of this acid is vital as it both maintains the acidic environment required for microbial function and provides reactants for the acidolysis-based dissolution pathways during the leaching process. As the biological growth curve (**Figure 1**) progresses, and the cultures reach the stationary phase, the metabolic pathways can be used to show peaked cell density through increasing Fe^3+^ concentration and biogenic acid. Accordingly, previous studies have tended to introduce battery electrode materials during the late exponential or stationary growth phases claiming maximized leaching activity at these phases.^31,32^ However, because these growth phases require long incubation times, the overall throughput of metal recovery can be constrained. The biological studies of ferric iron concentration [Fe^3+^], pH, oxidation-reduction potential (ORP), and biogenic [H_2_SO_4_], were monitored to gauge culture health and activity, with the metal’s recovery being determined via ICP-OES. Although redox-driven (redoxolysis) and acid-driven (acidolysis) mechanisms remain central to bioleaching behavior, the relationship between these parameters and actual metal dissolution is likely more complex and will require consideration with further work.

**Figure 1.**
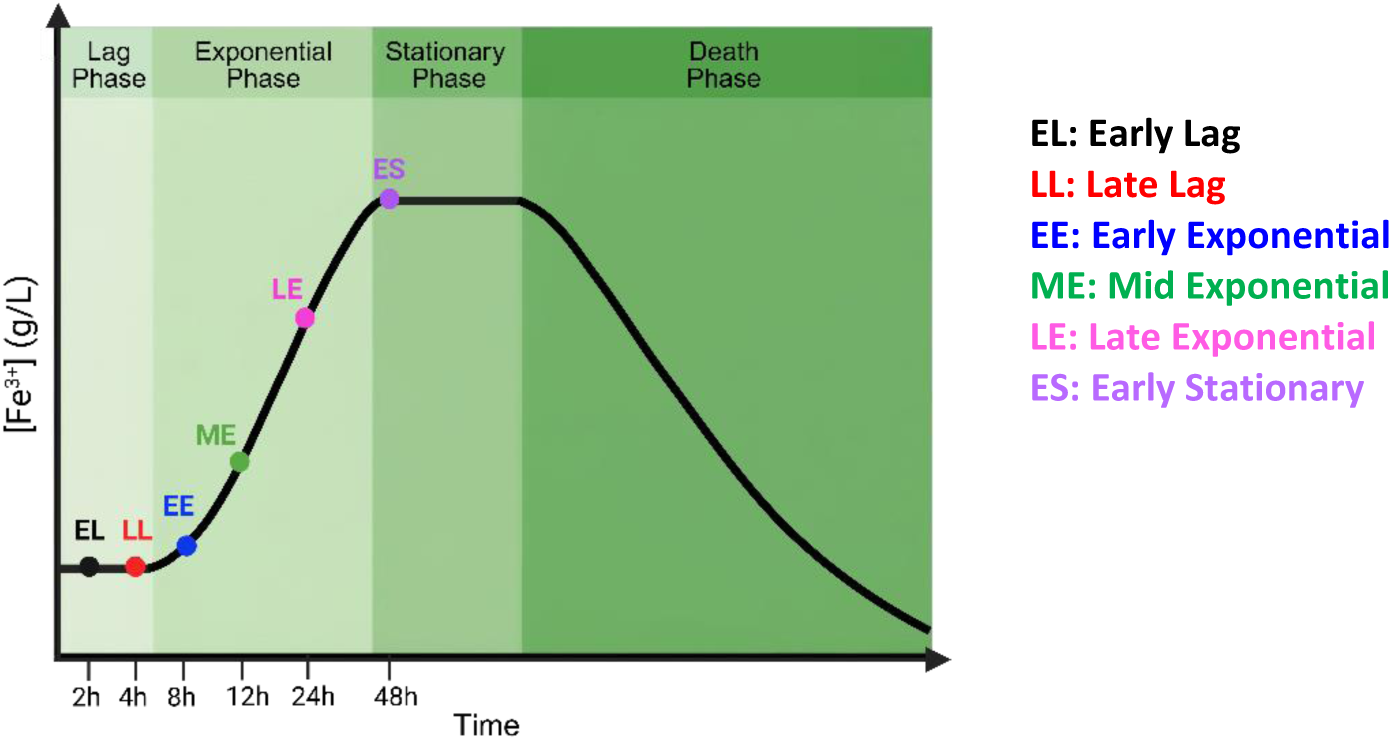
A schematic bacterial growth curve shows the major phases microorganisms go through during their lifecycle. The concentration of Fe^3+^ is used to represent cell growth due to the oxidation of Fe^2+^ during their metabolic cycle. The points along the curve represent the different incubation times *Atf* will undergo and where they will fall along the growth curve.

### 3.1. Biological Studies

For high-throughput studies, the experiments were first conducted in a 24 well-plate. The determined incubation period in this setup to reach stationary phase took between 48-72 hours and thus the incubation times of 2h (**EL**), 4h (**LL**), 8h (**EE**), 12h (**ME**), 24h (**LE**), and 48h (**ES**) were used to encompass the lag, exponential, and stationary phases. Using the EL incubation time as an example, the experiment proceeded as follows. The well plate was filled with M9K media and inoculated with a parent culture that had reached stationary phase. The initial 0h values of the four main biological data sets were taken. After 2h of incubation the biological data was taken again; at the same time, the NMC_622_ electrode materials were added to reach a pulp density of 10 g/L. Collection of the biological data was taken again 24h and 48h after the addition during the leaching period. The total time of incubation and leaching was 50 hours. After the leaching period the pregnant leaching liquor was filtered to remove any residual solid materials and then prepared for metals analysis using ICP-OES. This is repeated for all other incubation periods strictly differing only in the allowed initial incubation time prior to the NMC_622_ addition. In addition, supplementary trials both in biological growth studies (**Figure S1)** and the leaching efficiencies **(Figure S4)** are shown in the supporting information to indicate how different parent cultures may affect the growth and leaching trends. The overall conclusions can be drawn as the arching trends remain the same.

Monitoring the Fe^3+^ concentration in **Figure 2a** in the first 4 hours of incubation shows an initial moderate change from 3.97 ± 0.13 g/L at hour 0 to 4.15 ± 0.15 g/L and 4.14 ± 0.22 g/L, for EL and LL, respectively. This brief period of incubation shows the lag phase of the growth profile. From there they show more significant growth, reaching concentrations of 4.97 ± 0.18 g/L, 4.78 ± 0.22 g/L and 6.11 ± 0.27 g/L by the point of EE, ME, and LE, respectively, representing the exponential phase. Finally, at ES the Fe^3+^ reached a concentration of 8.69 ± 0.59 g/L showing the beginning of the stationary phases. When each of these incubation periods was reached, NMC_622_ electrode materials in a 10g/L pulp density were added to the culture and underwent leaching where the biological studies outlined herein were taken in 24 h increments for a total of 48 h. For the EL to ME based incubation cultures, [Fe^3+^] exhibited moderate changes in the first 24 h of leaching and remained consistent or slightly increased by the end of the 48 h of leaching. The result indicates that the attack of the Fe^3+^ ions on the NMC_622_ materials, aiding in their solubilization, was matched by the production of new Fe^3+^ from Fe^2+^ oxidation. The result strongly suggests that the cells did not experience a detrimental impact by the introduction of the cathode materials during early phases. However, in the systems incubated to LE and ES times, after the addition of NMC_622_, [Fe^3+^] decreased in the first 24 h of leaching and continued to go down by the end of the 48h of leaching. This can indicate that while the cathode material is being attacked for dissolution, *Atf* cells may be unable to continue growing, resulting in the depletion of [Fe^3+^].

**Figure 2.**
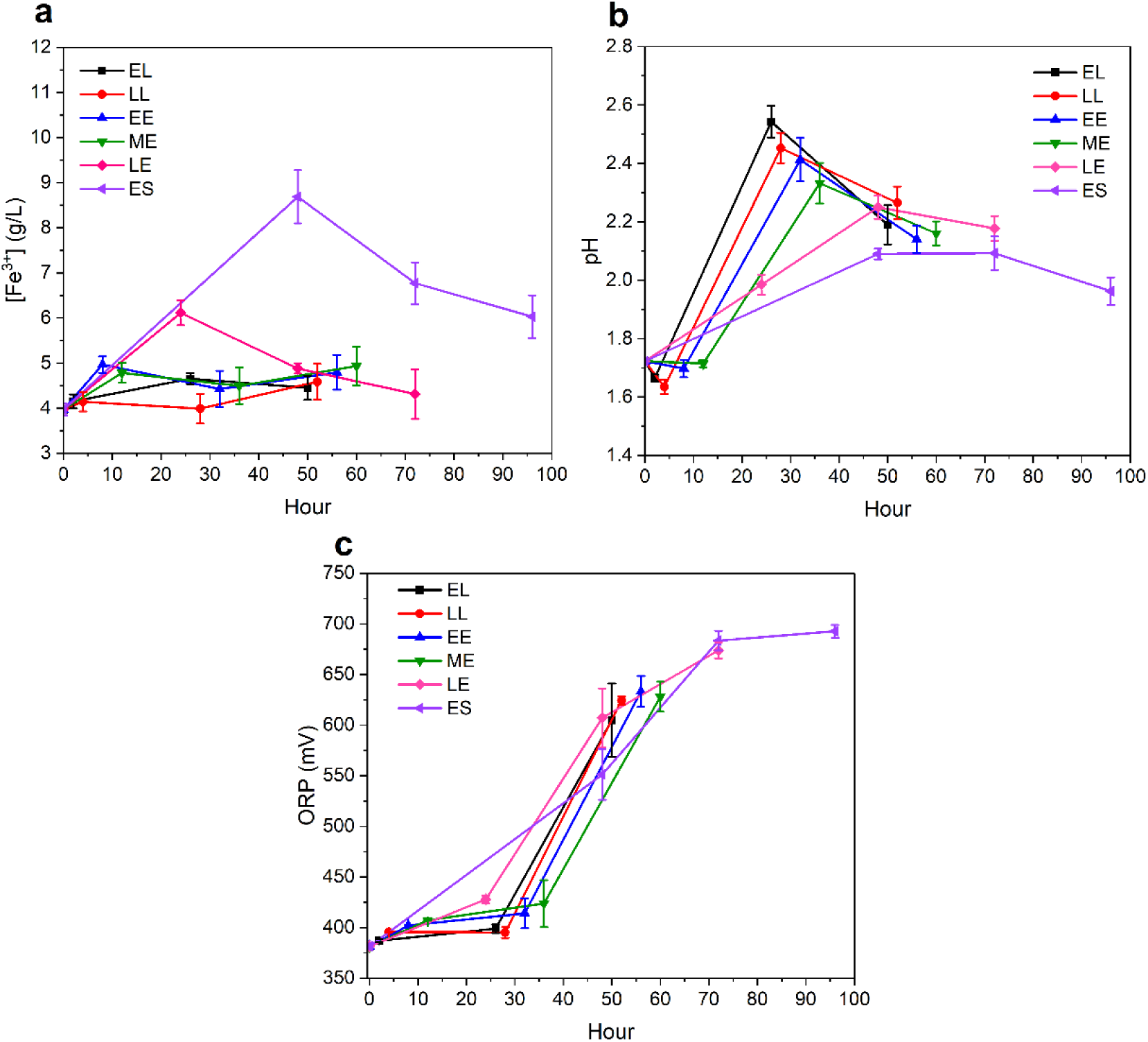
The biological studies of (a) [Fe^3+^], (b) pH and (c) ORP are used to indicate the cell growth and environment under incubation and leaching conditions. The cultures were each allowed to undergo incubation to specific time points of EL (black), LL (red), EE (blue), ME (green), LE (pink), and ES (purple) along with the resulting data during the leaching process where data was collected at 24 h intervals for 48 h total.

Along with [Fe^3+^], changes in pH during growth and leaching are important indicators of the effects NMC_622_ may impose when introduced to the culture at different growth phases. As shown in **Figure 2b**, all samples except ES exhibited a pH spike in the first 24h of leaching, followed by a decrease in pH in the next 24h, ending at a pH around 2.2. Notably, shorter incubation times were associated with larger pH variations. In contrast, the ES sample showed little to no pH increase during the first 24 hours of leaching and only a slight decrease by the end of 48 hours, reaching a final pH of 1.96 ± 0.05. This suggests that although higher biogenic acid production over time may buffer against pH spikes, the metabolic activity of the bacteria in the ES phase may not be sufficient to further lower the pH compared to early-stage cultures.

The third biological descriptor measured was the oxidation reduction potential (ORP), the data of which are presented in **Figure 2c**. ORP may be used as a measure on the Fe^3+^/Fe^2+^ ratio, and lower ORP values are associated with relatively lower [Fe^3+^] and vice versa. In the low incubation times of EL and LL, ORP underwent minimal changes compared to the initial value of 386 ± 0.8 mV, increasing slightly to 392 ± 2.2 and 396 ± 0.9, respectively. Further progressing along the growth curve to EE and ME stages, we measured modest changes to the ORP values of 405 ± 0.2 mV and 412 ± 0.3 mV, respectively. The result was also consistent with the nearly constant [Fe^3+^] as measured by spectrometer in this time window. As the growth continued from ME to LE and then ES, the ORP value increased steadily, showcasing continued oxidation of Fe^2+^ to Fe^3+^ as the culture approached peak metabolic activity. Referring back to **Figure 2a**, this trend coincided with the gradual rise in [Fe^3+^] for incubation times ME-ES where Fe^3+^ accumulated from *ca.* 4 g/L to 5-9 g/L. Notably, for cultures that experienced longer incubation times prior to the addition of NMC_622_ such as LE and ES, [Fe^3+^] deceased sharply in the first 24 h of leaching before further declining by the end of the 48 h leaching period. In contrast, the ORP continued to rise or plateaued at these same time points. Upon addition of NMC_622_ for leaching, all ORPs shifted to higher absolute values over the same time frame, while [Fe^3+^] remains relatively stable or even decreased for some conditions, further underscoring that ORP captures the integrated electrochemical response of the system rather than simply tracking bulk [Fe^3+^].

It is noted that ORP and [Fe^3+^] are not expected to act identically over the course of leaching. This is because the former reports on the mixed electrochemical response of redox active species at the electrode interface, whereas the latter reports on the bulk, operationally defined pool of Fe^3+^. In practice, the potential recorded by an ORP probe is governed primarily by the local activities of Fe^3+^ and Fe^2+^ near the electrode surface, which can diverge from the total Fe^3+^ concentration because complexation with sulfate and transition metals, as well as precipitation into phases such as jarosite or schwertmannite (an Fe(III) oxyhydroxy-sulfate), can remove Fe^3+^ from the electrochemically accessible pool without strongly changing the bulk analytical signal. Additional redox couples introduced during NMC_622_ dissolution, as previously mentioned, may further contribute to the mixed potential and can drive substantial ORP shifts during the first 24 h of leaching even when the spectrophotometrically determined Fe^3+^ concentration changes only slightly. The formal Fe^3+^/Fe^2+^ potential in sulfate media is also sensitive to pH and speciation. As such, proton consumption by NMC_622_ and subsequent acid generation by *Atf* metabolism can alter ORP at essentially constant bulk [Fe^3+^].

Finally, as acidolysis is a key factor for high leaching efficiencies of desired metals, the biogenically produced acids will aid in the dissolution of metals by following the reactions below^33^:

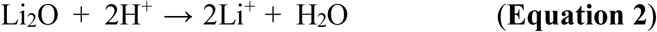

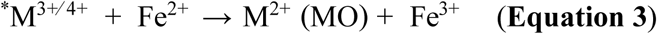

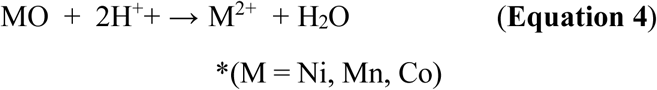

To determine the acid concentrations at each incubation timepoint, acid-base titration was taken, and the data are shown in **Figure 3**. The biogenic H_2_SO_4_ concentration remained relatively low in the first 12 h of incubation, reaching 0.017 ± 0.003 M, 0.011± 0.001 M, 0.015 ± 0.002 M, and 0.026 ± 0.007 M for EL, LL, EE, and ME, respectively. This correlated well with [Fe^3+^] that showed there was no significant cell growth as it was in the early stages of the biological cell growth curve. However, by incubation stages of LE and ES, the acid concentrations reached 0.135 ± 0.021 M and 0.401 ± 0.036 M, respectively. Previously literatures have stated that the most efficient battery cathode leaching occurred when acid concentrations were between 0.17-0.52 M.^34^ This was thought to be beneficial when compared to the much higher acid concentrations (1-4 M) required in the hydrometallurgy approach,^35–37^ which present significant challenges for downstream waste water management. In our systematic study, we observed that there was sufficient leaching in cultures early in their growth stages. The result further indicated that low acid molarity was sufficient for metal solubility by *Atf*, suggesting a process that does not simply depend on acidolysis.

**Figure 3.**
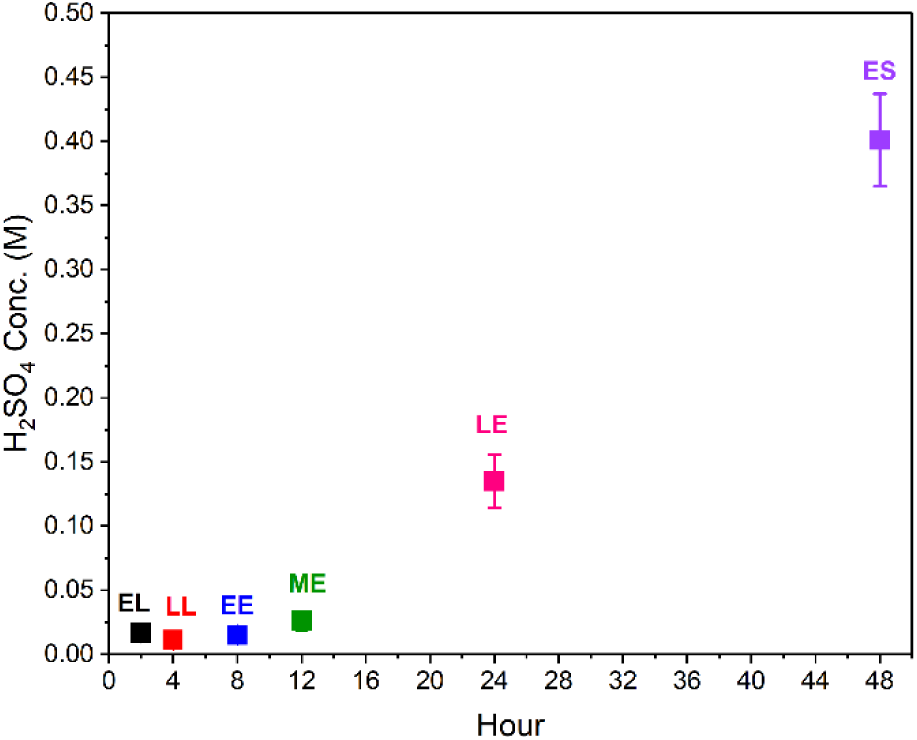
The measured biogenic H_2_SO_4_ concentrations at different *Atf* growth stages show the physiological state of the specific incubation times. As the cultures are allowed to mature, the [H_2_SO_4_] increase as the reduced sulfur species in the media interact with produced protons by the metabolism of *Atf*.

**Table 2.**
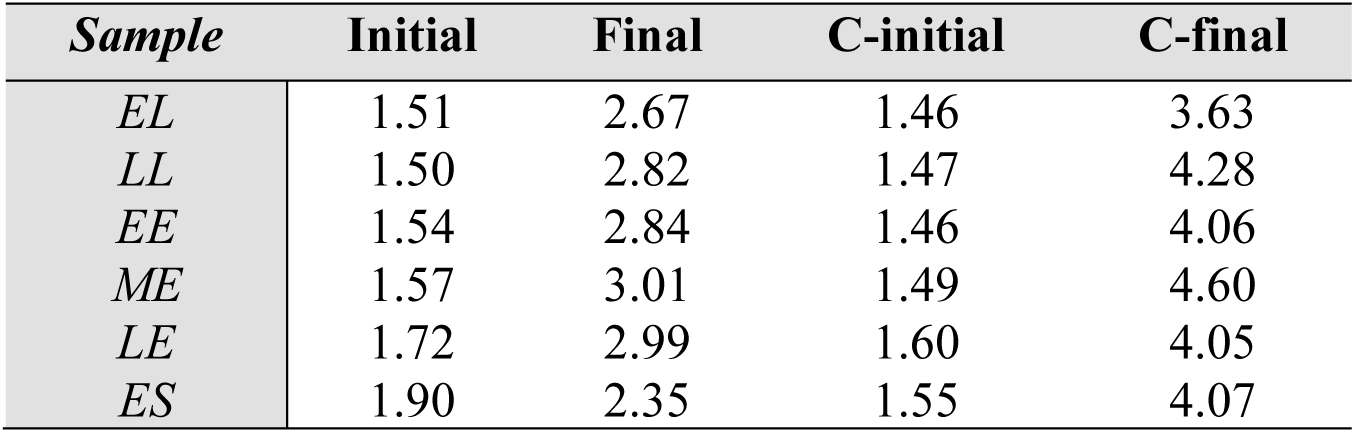
The initial and final pH values of the culture systems. The final pH is after 48 hours of leaching. The initial and final concentrations of the control systems show the regenerative capacities of the *Atf* containing leaching systems. C-initial and C-final indicate control cases, in which no cells were introduced.

### 3.2. Leaching Efficiencies

When the cultures had reached their intended incubation times, the cathode powder was introduced for direct contact leaching. Li for all time points showed the highest leaching efficiency, likely due to its readily solubilized nature in that acidic environment. However, notably, incubation points of EL-LE showed comparable leaching efficiencies across all metals (**Figure 4**). This differed drastically for 48-hour incubation time, which should put the culture in early stationary phase, with a significant drop off in leached metals. Since the pH remained low for the 48h incubated culture prior to leaching suggests that the bioleaching process is more complex than simple acidosis. One explanation is that there is an imbalance of physiology and chemistry. In this view, “acid availability” and “leaching machinery” (i.e., the Fe^3+^ reserve, redox capacity, and active cells) both are required, rather than a simple pH effect, for efficient leaching. When using low incubation time cultures (e.g., EL at 2h) while there is less total biogenic acid, there may be a more optimal combination of active cells and Fe^2+^/Fe^3+^ chemistry for the attack on NMC_622_ compared to that of the long incubation time (e.g., ES at 48h). This happens despite the bulk pH of EL spiking higher to a pH of 2.54 versus a pH of 2.09 in the ES case.

**Figure 4.**
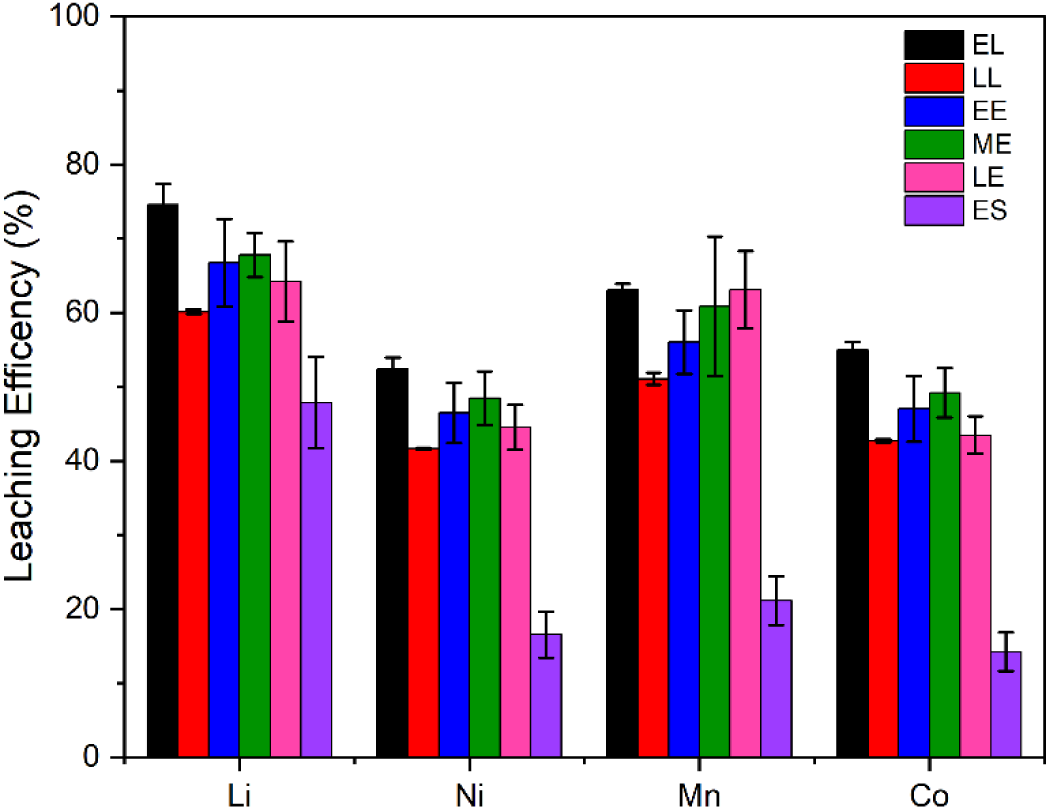
The leaching efficiency of metals recovered from model electrode (NMC_622_) powder using a traditional bioleaching method of Atf (M9K-FS) that is allowed to incubate for specific time frames. These leaching efficiencies show the systematic study of how incubation times can affect metals recovery.

In ES, several factors may impact the bioleaching performance. The *Atf* cells in the stationary phase exhibit slower metabolic activities, entering a “maintenance” mode. This may result in a slower Fe^2+^ oxidation and lower acid generation per cell compared to cells that are in EL or even EE. Additionally, due to the aged media in ES, Fe^2+^ is already largely oxidized to Fe^3+^ with some Fe^3+^ lost due to the formation of jarosite, seen as the orange precipitate formed at pH values 1.5-2.5.^38,39^ When in large quantities, these precipitates could passivate the NMC_622_ powder surface and consume protons leading to decrease leaching performance.^40^ When NMC_622_ is added to the culture at a later stage, there is not sufficient active Fe^2+^/ Fe^3+^cycling to maintain a high, sustained oxidizing environment on the particle surface during direct contact leaching. A previous work on LiCoO_2_ and mixed cathodes recovery also noted that metal toxicity plus insufficient Fe^3+^ regeneration collapsed leaching at higher pulp densities or stressed cultures, even when pH was acceptable.^41^ In short, while the stationary culture may seem to have acceptable pH if there is poor dynamic capacity to keep generating Fe^3+^ and protons, then leaching of NMC_622_ would be limited.

The difference in leaching performance between ES and EL can be attributed to the physiological state of the cells. In the early lag phase (EL), cells are transitioning to a highly metabolically active state and, unlike stationary phase cells, are not yet nutrient-limited. Consequently, specific Fe^2+^ oxidation rates are high.^42^ At the time of NMC_622_ addition, due to the early incubation nature, there is usually a larger pool of Fe^2+^ present in the medium. Thus, when NMC_622_ is added, *Atf* can rapidly regenerate Fe^3+^ and acid, creating a strong indirect leaching system even if the initial bulk pH jumps to ∼2.7 from NMC acid consumption or alkali release. This supports our hypothesis that early exposure to NMC_622_ may also allow cells to adapt gradually to dissolved Ni, Co, and Mn, maintaining activity rather than encountering these metals after becoming stressed and stationary.^43^ As a result, more effective Fe^3+^ cycling and proton production occur over the first 24–48 hours of leaching, so even with a resulting final pH of 2.7, the cumulative oxidizing/acidic attack on NMC is stronger, resulting in roughly double the leaching efficiency.

While the 24-well plate format was used for high-throughput systematic studies, we recognize that increasing the scale may alter the leaching efficiency. To examine this effect, we repeated the experiment in standard test tubes at a volume of 4 mL (3.6 mL media and 0.4 mL inoculant), with selected incubation times: EL, EE, LE, and ES. The biological measurements of [Fe^3+^], pH, and ORP (**Figure S3**) showed an overall trend like the 24-well plate experiments for the EL-ES incubations. Importantly, the resulting leaching efficiencies (**Figure 5**) showed enhanced metal recovery across all phases compared to 24-well plates. In this system, the LE and importantly the ES incubations showed the highest metals leached with LE reaching 89% (Li), 92% (Ni), 92% (Mn), and 93% (Co) and ES reaching 94% (Li), 96% (Ni), 96% (Mn), and 96% (Co). The EL phase, incubated only for 2 hours, was still capable of leaching at comparable efficiencies reaching 86% (Li), 86% (Ni), 86% (Mn), and 86% (Co) in the same 48-hour period. This finding demonstrates that a two-step bioleaching process, where the cultivation is brief before introducing NMC_622_ electrode materials, is a viable option and may address concerns regarding the slow kinetics of bioleaching.

**Figure 5.**
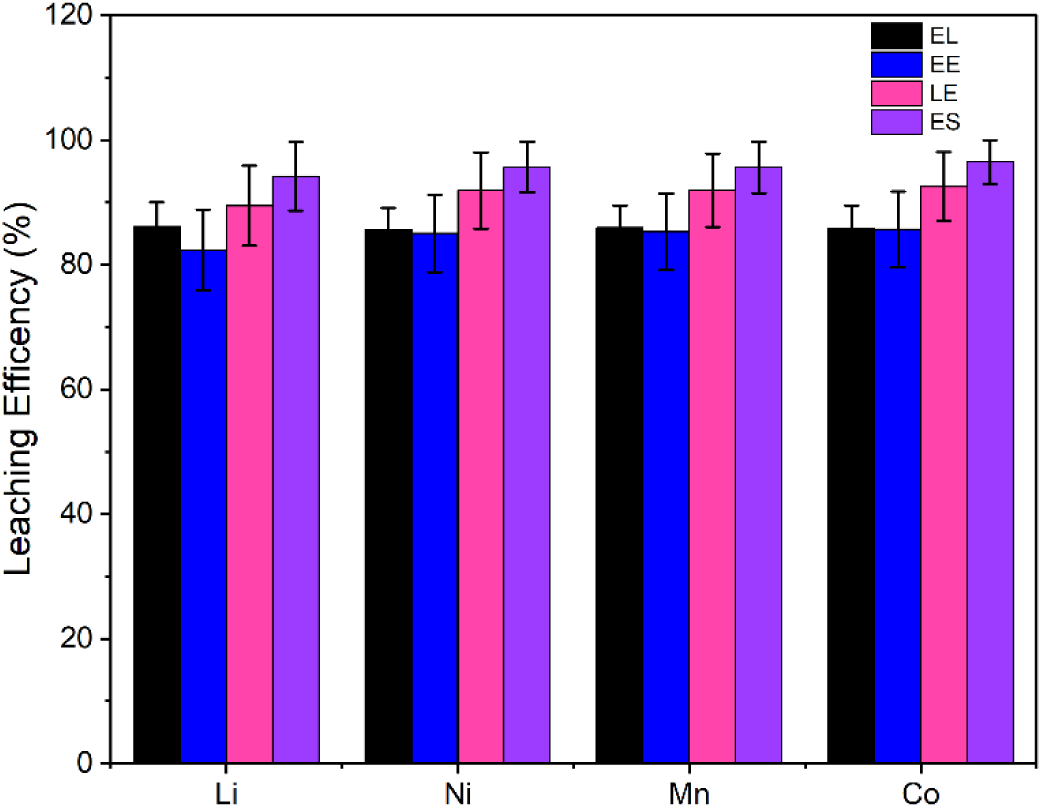
The leaching efficiency of metals recovered from model electrode (NMC_622_) in scaled up test tube systems. The culture incubation times studied were EL (2h, black), EE (8h, blue), LE (24h, pink), and ES (48h, purple). The leaching efficiencies show near similar recovery rates regardless of incubation time.

Beyond the factors of the culture, we see that in leaching efficiency studies performed in test tubes, the overall leaching efficiency was compared to that of our previously reported values.^44^ Even though the leaching efficiencies were lower in the 24-well plates, the overall trends remained the same (**Figure 5**). While cultures with longer incubation periods showed the highest [Fe^3+^], lowest pH, and highest ORP, they achieved the lowest leaching efficiencies for the metals of interest in the 24-well plate format. This may be due to the culture reaching a point of being effectively “over-oxidized” and passivated rather than actively dissolving the solid. At very high ORP and ferric levels, the excess Fe^3+^ tends to form surface precipitates such as jarosite or schwertmannite that can coat the NMC_622_ particles, blocking the surface from the acid and oxidants. In sulfide rich systems, this effect is well documented: once the optimal window is exceeded, leaching rates drop sharply and the media minerals enter a passive state.^45^ Under these conditions, the solution may look to be strongly oxidizing by the biological growth metrics, but the reaction taking place at the solid to liquid interface could be transport-limited. As a result, the chemical attack of Fe^3+^ and H^+^ on the cathode material becomes sluggish and less efficient, with the added effect that excessively low pH and high Fe^3+^ can stress *Atf* and shift its metabolism toward maintaining redox balance and proton gradients rather than supporting further Fe^3+^ regeneration and biofilm formation on the particles, which promote leaching. Consequently, the bacterial “catalyst” becomes less effective despite favorable seemingly chemistry. Finally, if later-stage cultures with the most extreme [Fe^3+^]/ORP conditions also experienced stronger metal toxicity or nutrient limitations, the resulting system features high oxidizing potential but poor mass transfer and compromised biocatalysis. This trade-off can balance the overall leaching performance compared to cultures operating in more moderate redox conditions.

### 3.3. Mathematical Model

To examine how different parameters contribute to the dynamics of bioleaching, we constructed a simplified mathematical model that describes the process of bioleaching. This model is based on a simple assumption that bacteria are producing an agent (which could be Fe^3+^ or biogenic acid) that moves the bioleaching process forward. Based on this assumption, we simulate the dynamics of *Atf* growth and leaching. To simulate the growth of *Atf*, we assume that bacteria have a lag time of *t_l_*, follow logistic growth (with a maximum growth rate of *r* and a carrying capacity of *K*), and are inhibited by metal toxicity from the presence of NMC_622_ material. Thus,

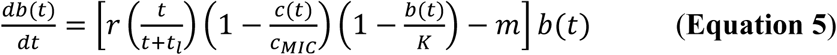

where *b*(*t*) is the bacterial cell density as a function of time, *c*(*t*) is the concentration of NMC_622_ as a function of time, *c*_*MIC*_is the minimum inhibitory concentration of NMC_622_ (i.e., the lowest NMC_622_ concentration at which bacteria can no longer grow), and *m* is the mortality rate of bacteria. To model the concentration of NMC_622_ over time, we use the following equation:

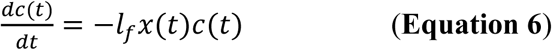

where *x*(*t*) represents the leaching agent produced by bacteria (presented in arbitrary units of U). We assume that *x* is produced at a steady rate *p* by bacteria and gets removed with the same rate that NMC_622_ is converted (proportional to *dc*(*t*)⁄*dt*, with a consumption factor *s*). A leaching factor, *l*_*f*_, represents how well the leaching agent converts the NMC_622_ material into the leaching products.

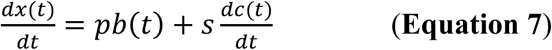

**Figure 6** shows the simulation results, using the parameter values listed in **Table 3**. **Figure 6a** shows the growth dynamics of *Atf*. A close examination of **Figure 6a** shows a decrease in growth rate at the time of introducing the NMC_622_ material into the culture, consistent with the assumption of such inhibition in the model. **Figure 6b** shows how NMC_622_ is converted in the process of bioleaching. The initial spike marks the time when the NMC_622_ material is added in, and bacterial activity converts this material over time through the process of bioleaching. Noticeably, the rate of conversion is smaller when the NMC material is added earlier (and the bacterial cell density and the density of the bioleaching agent is lower). This rate increases if the material is added after a longer initial incubation time. Nevertheless, the simulated dynamics shows that in all cases, the majority of the cathode material can be converted, reaching a high overall leaching efficiency. **Figure 6c** shows the concentration of the leaching agent, which grows as the bacterial cell density increases. This value is slightly lower in cases where the NMC_622_ material is added earlier, because a portion of the leaching agent is participating in leaching the material and is consumed earlier. **Figure 6d** compares the final leaching efficiencies at varying NMC_622_ introduction times that match our experimental protocol. In all these cases, the leaching efficiency is calculated after 48 hours of leaching following the introduction of the NMC_622_ material. The trend in **Figure 6d** qualitatively matches that of the scaled-up experiments in **Figure 5**.

**Figure 6.**
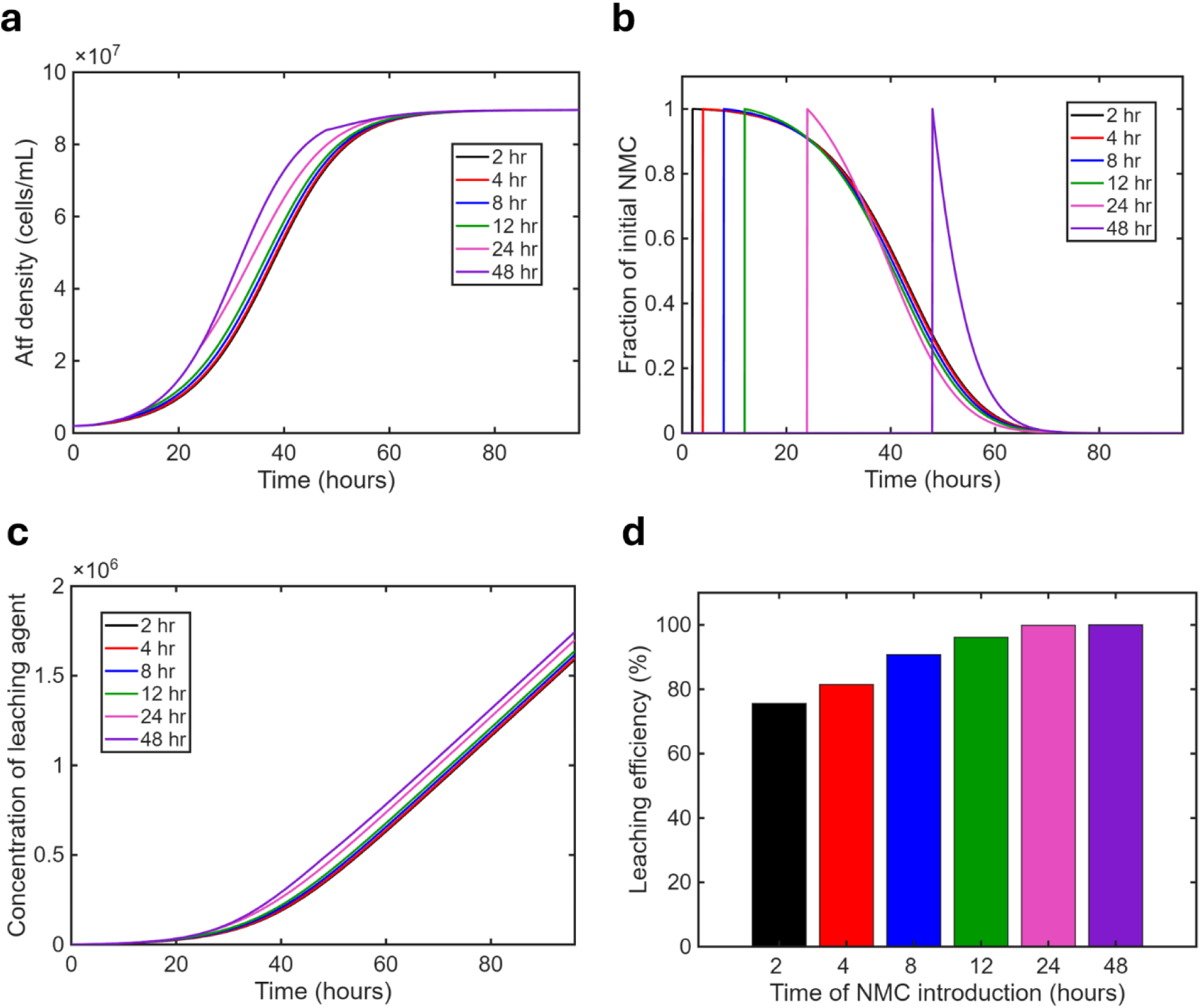
Simulated dynamics of bioleaching creates predictions comparable to experimentally observed results. Simulated values of (a) bacterial population density, (b) remaining NMC_622_, (c) the concentration of the leaching agent, and (d) the overall leaching efficiency after 48 hours of leaching are shown. Equations 5-7 are used for these simulations, with parameters listed in Table 3.

**Table 3.1.** The description and values of the parameters of the model used for simulating the bioleaching dynamics are listed.

| Parameter | Description | Value |
| --- | --- | --- |
| $r$ | *Atf* growth rate | $0.2 \text{ h}^{-1}$ |
| $K$ | *Atf* carrying capacity | $10^8 \text{ cells/mL}$ |
| $m$ | *Atf* death rate | $0.02 \text{ h}^{-1}$ |
| $l_f$ | Leaching factor | $10^{-6} \text{ U}^{-1} \cdot \text{h}^{-1}$ |
| $p$ | Leaching agent production rate | $3 \times 10^{-4} \text{ U}/(\text{cell} \cdot \text{h})$ |
| $s$ | Leaching agent consumption rate | $10^4 \text{ U/mg}$ |
| $C_{MIC}$ | Minimum inhibitory concentration of metal toxicity from NMC | $50 \text{ g/L}$ |
| $b_0$ | Initial *Atf* concentration | $2 \times 10^6 \text{ cells/mL}$ |
| $C_0$ | Initial NMC cathode material concentration | $10 \text{ g/L}$ |

We explored how different parameters could affect the leaching efficiency; we simulated the dynamics by varying one parameter at a time. We first examined how the strength of inhibition of bacteria by NMC_622_ affected the leaching efficiencies. As shown in **Figure S6**, stronger NMC inhibition leads to lower leaching performance in cultures to which NMC_622_ is added after shorter incubation times. This is because earlier cultures are initially slower in their leaching activity, thus the bacterial culture spends more time under stronger inhibition by NMC_622_, grows more slowly under such condition, and overall shows less efficiency during the fixed 48-hour leaching. With weak NMC_622_ inhibition, the difference between early versus late introduction of NMC_622_ was less, whereas this difference was magnified when NMC_622_ inhibition was stronger.

We then investigated how the production rate of the leaching agent by bacterial cells affected the leaching efficiencies. As shown in **Figure S7**, lower rate of production of the leaching agent leads to inferior leaching performance in cultures to which NMC_622_ is added after shorter incubation times. This is consistent with the results in **Figure S6**, because less production of the leaching agent still leads to the bacterial culture spending more time under inhibition by NMC_622_, and thus a lower overall efficiency. When the leaching agent is produced in abundance, the difference between early versus late introduction of NMC_622_ becomes less pronounced. We observed the same trends when the consumption rate of the leaching agent was changed: A lower rate of leaching agent consumption diminished the difference between early versus late introduction of NMC_622_, whereas a higher rate of leaching agent consumption increased that difference (**Figure S8**). Notably, this effect is weaker, even with order-of-magnitude changes in consumption rates.

The main takeaway from our simulations is that if the leaching performance was weak (e.g., because of the rate of production of the leaching agent being small) or if there was strong inhibition of *Atf* by the NMC_622_ material, we predict to see the early-lag case significantly underperforming in leaching efficiencies. This was not the case based on our experimental observations, suggesting that the intermediate values presented in **Figure 6** offer a reasonable estimate of the relative strength of different factors (particularly, NMC_622_ inhibition and leaching performance). We also note that this simple model does not match the observations in **Figure 4** with EL case overperforming in leaching efficiencies, suggesting that in such cultures there is an additional process in effect which is not accounted for in our simple model. We defer a more thorough exploration of the potential cause to future investigations.

## 4. Conclusions

This work aims to systematically test the ability of *Atf* to extract metals from NMC_622_ electrode powder when cells are in different growth stages. By doing so we provide insights into how leaching efficiency is affected by cell physiology and biogenic acid production. We highlight how, unlike previously thought, the wait to reach exponential or stationary phases is not necessary for metals recovery. When NMC_622_ is added during an early incubation phase (e.g., EL, 2h), leaching efficiencies are comparable to or even exceed those observed at later phases. Furthermore, our scaled-up experiments show the same trend, with improved leaching efficiencies aligning with our previously reported work. Specifically, after only 2 hours of incubation (early lag phase) and 48 hours of leaching, efficiencies of 59% (Li), 31% (Ni), 40% (Mn), and 31% (Co) were achieved. In comparison, 48h of incubation time (stationary phase) and 48 hours of leaching resulted in 48% (Li), 17% (Ni), 22% (Mn), and 16% (Co) recovery. Thus, initiating leaching during the early incubation phase of *Atf* enables higher metal recovery in approximately half the total time. Efficient bioleaching using early culture stages reduces the need for prolonged cultivation period and can reduce the logistic burden of LIB recycling via BioHM.

Future work will investigate the genomic profiles of these cultures at the designated time points to determine if certain variants are selected when the NMC_622_ material is added to the early stages of the culturing process. Additionally, the study of how cultures at different incubation times respond to increasing pulp densities will reveal the limits of early phase cultures for leaching. With these studies accomplished, evolutionary insights may be applied to counter the slow kinetics bottleneck and improve overall performance of the biohydrometallurgy method.

## Author contributions

Brooke. E. Elander; formal analyses, methodology, investigation, data curation, conceptualization, writing-original draft preparation; Mengyun Jiang: investigation, methodology, investigation, writing-original draft preparation; Cameron Guthrie and Zachary Ibrahim: investigation, writing-review; Babak Momeni and Dunwei Wang: conceptualization, supervision, review, and editing.

## Conflicts of interest

There are no conflicts to declare.

## Data availability

Data will be made available upon request.

## Acknowledgements

The authors would like to acknowledge Undergraduate Research Fellowships from Boston College that supported MJ and CG. This work was supported by the National Science Foundation (NSF CET EAGER) under Grant No. 2342967.

## Supporting Information

### 1. Growth profile of *Atf*

#### 1.1 Various trials in 24-well plate system

The experiments were conducted in a 24-well plate (Fisher scientific) setting with 1.5mL of M9K media in each sample. All samples’ pH were adjusted to 1.6-1.8 using 2M H_2_SO_4_ (Millipore Sigma) for the initial acidification step. Each sample was inoculated with 10% v/v parent culture and went through incubation at 30 °C under shaking at 170 rpm. The 24 wells were divided into 6 groups: EL (2h), LL (4h), EE (8h), ME (12h), LE (24h), and (48h), with 4 replicates each. This process was replicated across multiple trials. What is a keynote is that due to the usage of different parent cultures being used as the inoculant they exact features of biological growth and the leaching efficiencies may vary. However, the trend of the data does remain the same. As seen in **Figure S1a-c.**

**Figure S1.**
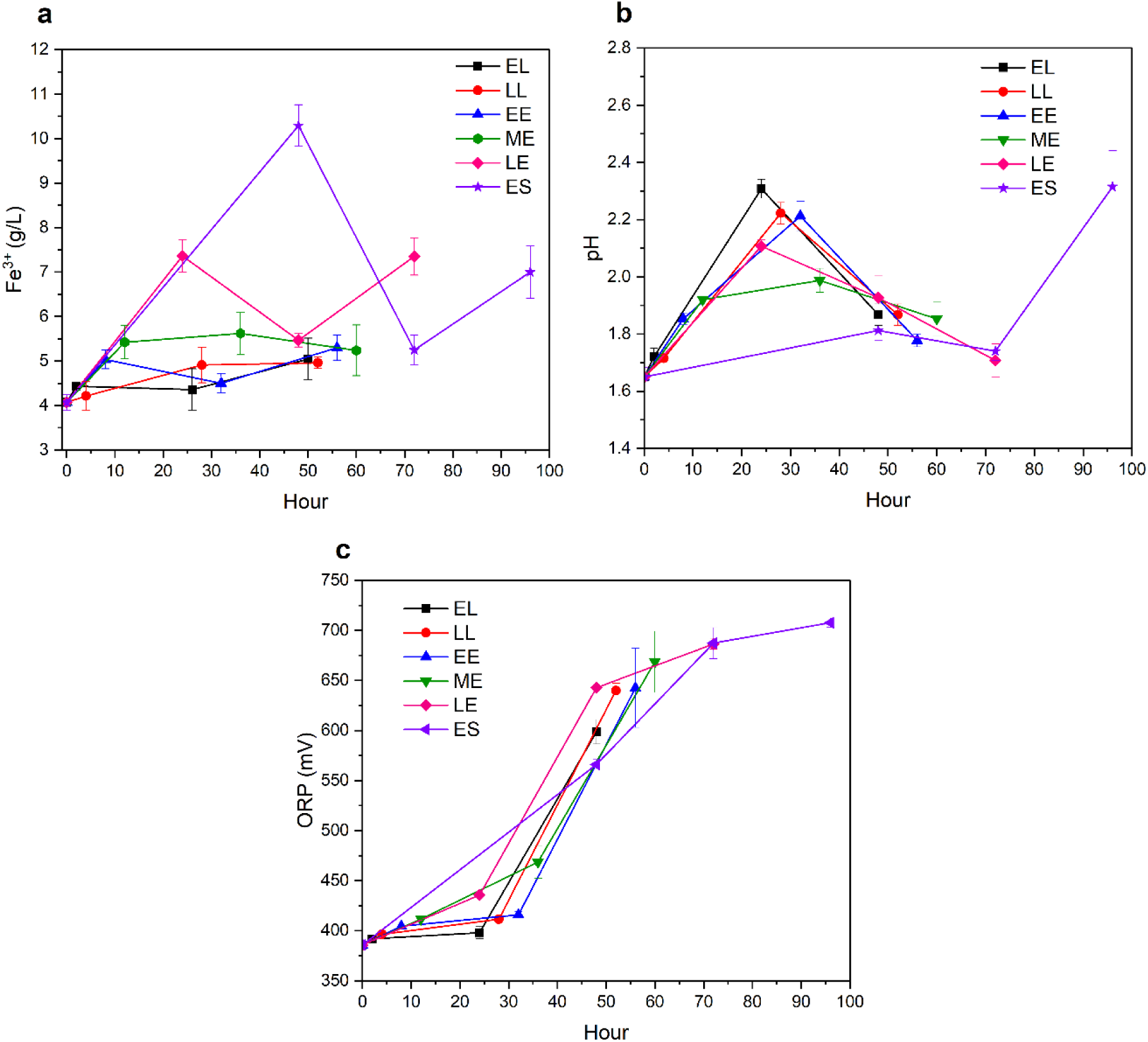
The (a) [Fe^3+^], (b) pH, and (c) ORP profiles of other trial for the timed incubation versus leaching efficiencies.

#### 1.2 Control trial

The control experiments, **Figure S2**, were also conducted in a 24-well plate (Fisher scientific) setting with 1.5mL of M9K media in each sample. All samples’ pH were adjusted to 1.6-1.8 using 2M H_2_SO_4_ (Millipore Sigma) for the initial acidification step, however there was no addition of *Atf.* The control wells still went through the same steps as the inoculated wells and were under incubation at 30 °C with shaking at 170 rpm.

**Figure S2.**
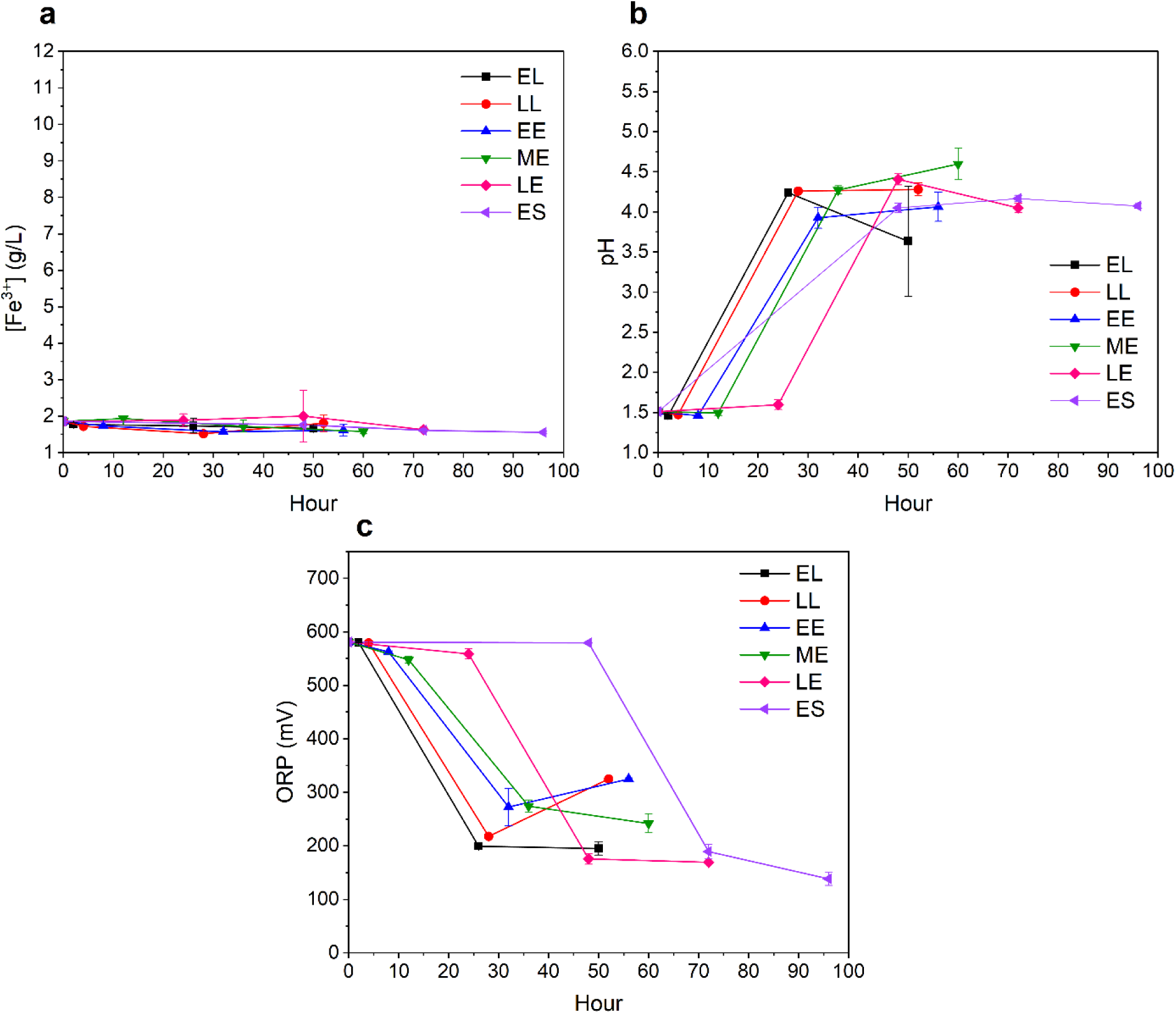
The (a) [Fe^3+^], (b) pH, and (c) ORP profiles of the controlled systems for the timed incubation vs leaching efficiencies

#### 1.3 Test-tube scale

**Figure S3.**
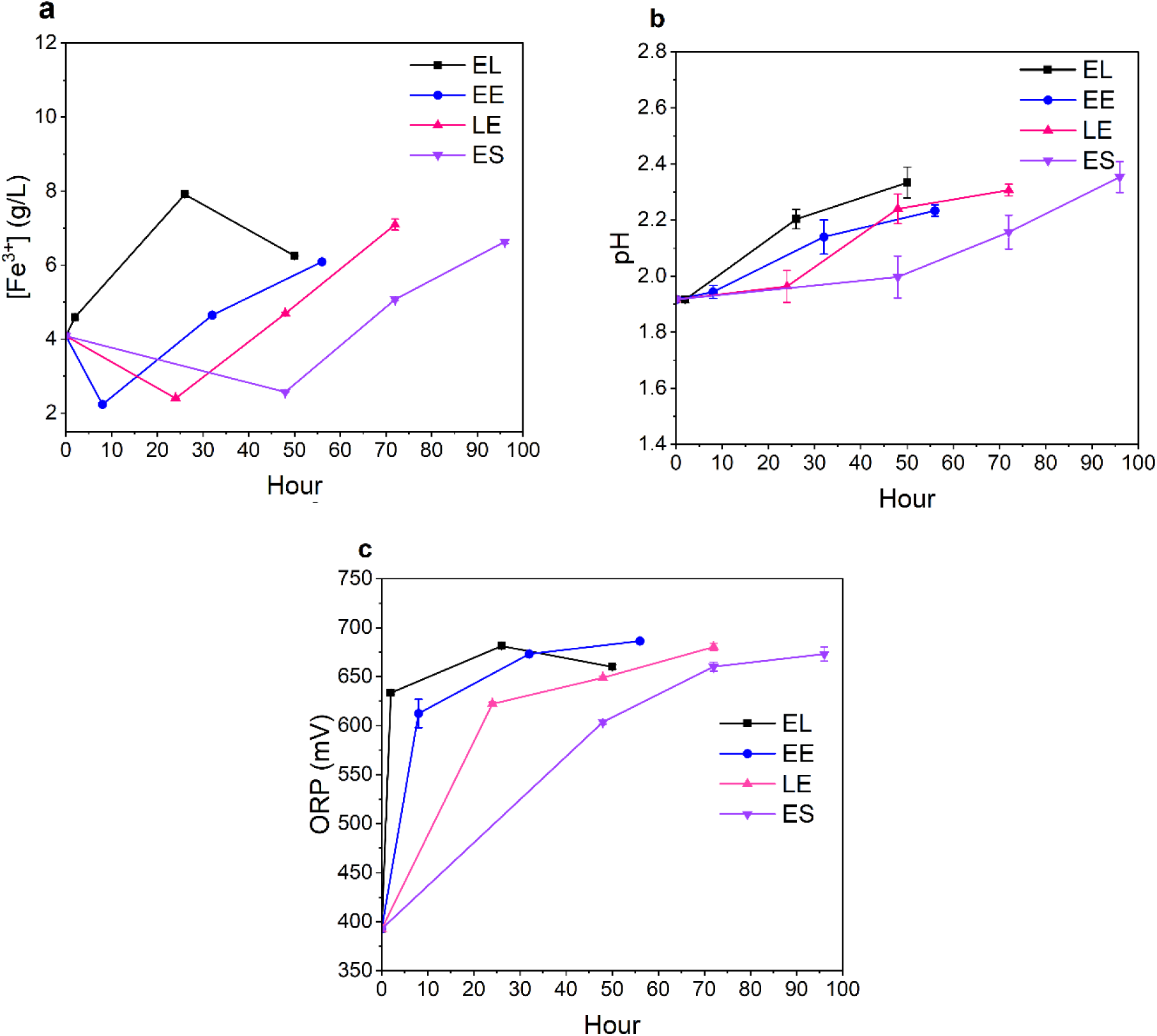
The (a) [Fe^3+^], (b) pH, and (c) ORP profiles of the test tube scaled up trial.

### 2. Leaching efficiencies under varied systems

#### 2.1 24-well plate various trials leaching efficiency

The trial two of the timed culture growths leaching efficiencies show how slight differences in parent cultures may lead to slight variations in the exact values. This may be due to the deviations that occur from culture to culture which can come from variations in culture health, mutations or other cell level changes. What is noted however is that across both trials the additions of NMC_622_ to the low incubation times showed enhanced leaching efficiencies compared to that of later stage incubations

**Figure S4.**
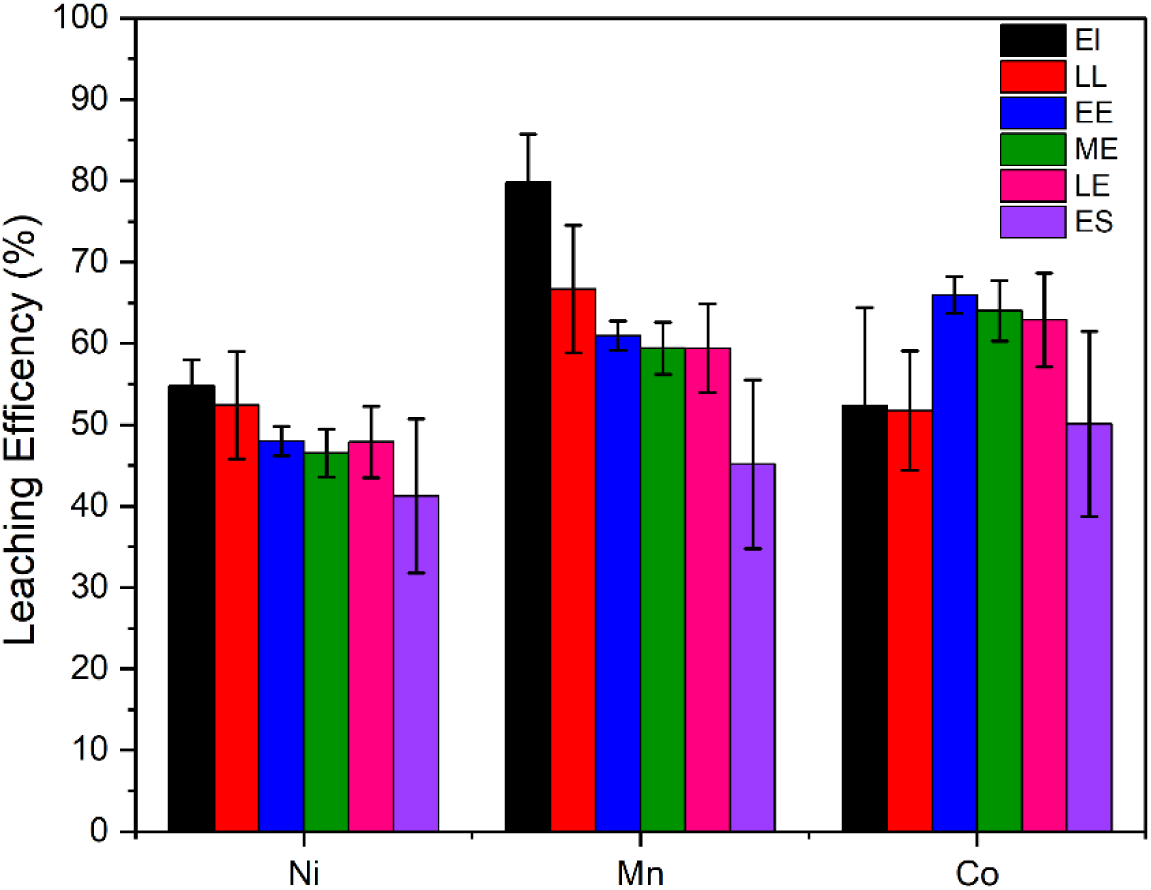
The leaching efficiencies in a trial of the 24 well plate setting whose biological growth data is outlines in Figure S1.

#### 2.2 Control in 24-well plate

**Figure S5.**
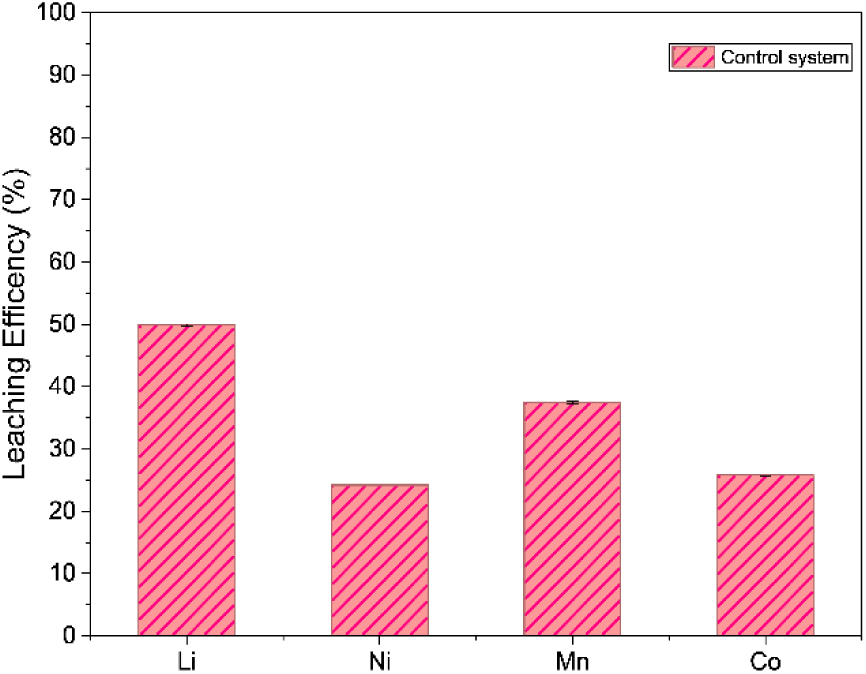
The leaching efficiency in the control 24 well plate setting whose biological data is outlines in Figure S2.

### 3. Modeling leaching efficiencies under alternative parameters

We investigated the contributions of different processes by modulating their corresponding parameters in the model presented in **Equations 5-7**.

**Figure S6.**
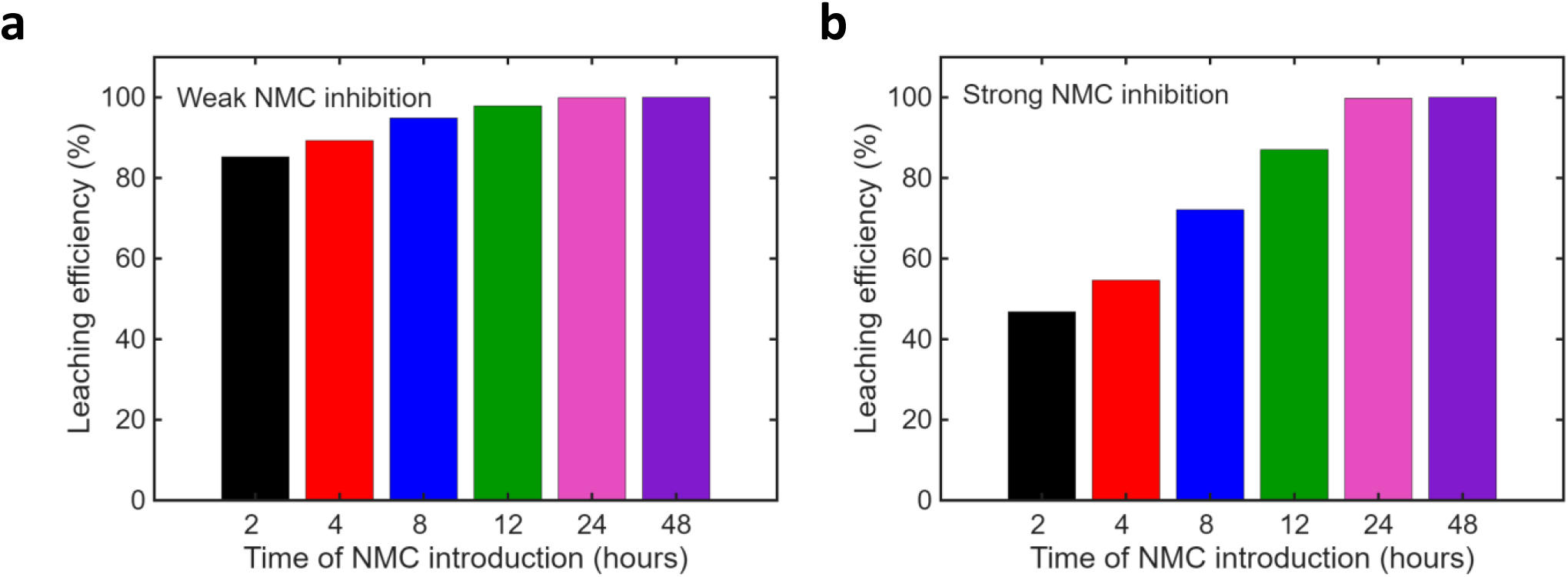
Simulated leaching efficiency after 48 hours of leaching is shown for cultures initiated with the same cell density but with different times of NMC introduction. All the parameters match the values listed in Table 3, except the minimum inhibitory concentration of metal toxicity from NMC. (a) Weak NMC inhibition, *c*_*MIC*_ = 25 g/L. (b) Strong NMC inhibition, *c*_*MIC*_ = 2 g/L.

**Figure S7.**
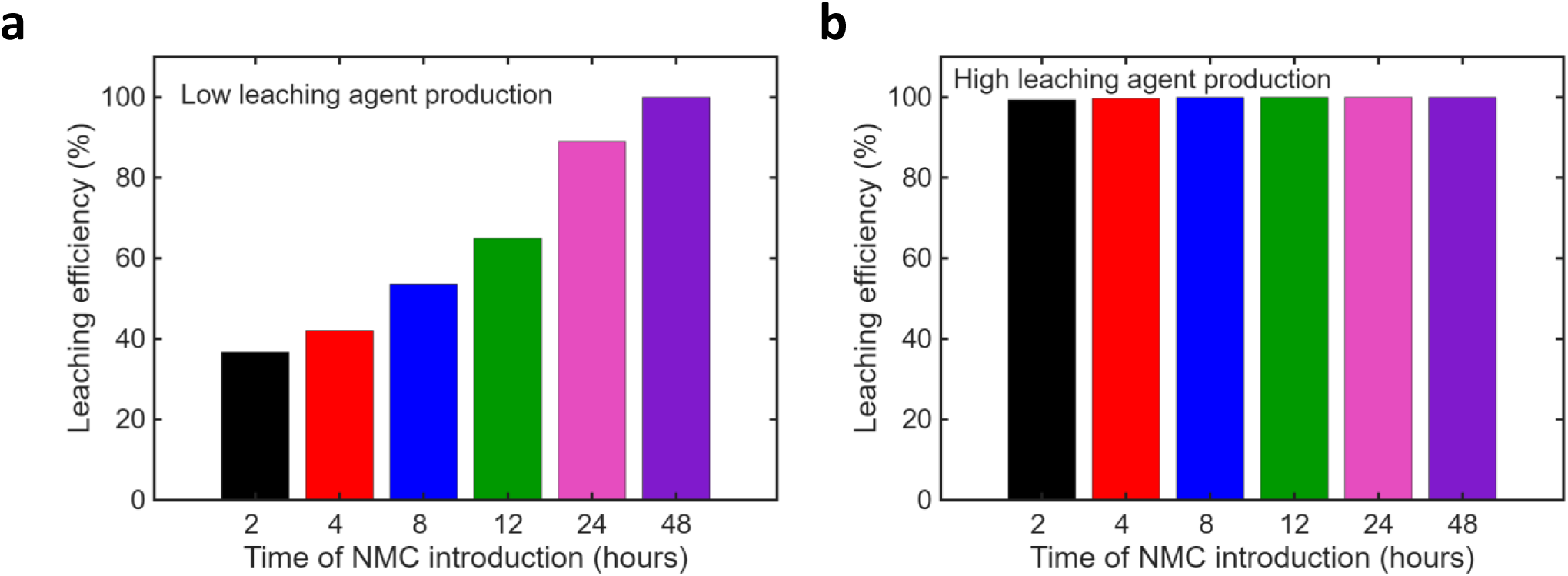
Simulated leaching efficiency after 48 hours of leaching is shown for cultures initiated with the same cell density but with different times of NMC introduction. All the parameters match the values listed in Table 3, except the bacterial production rate of the leaching agent. (a) Low leaching agent production rate, *p* = 10^-4^ U/(cell‧h). (b) Low leaching agent production rate, *p* = 10^-3^ U/(cell‧h).

**Figure S8.**
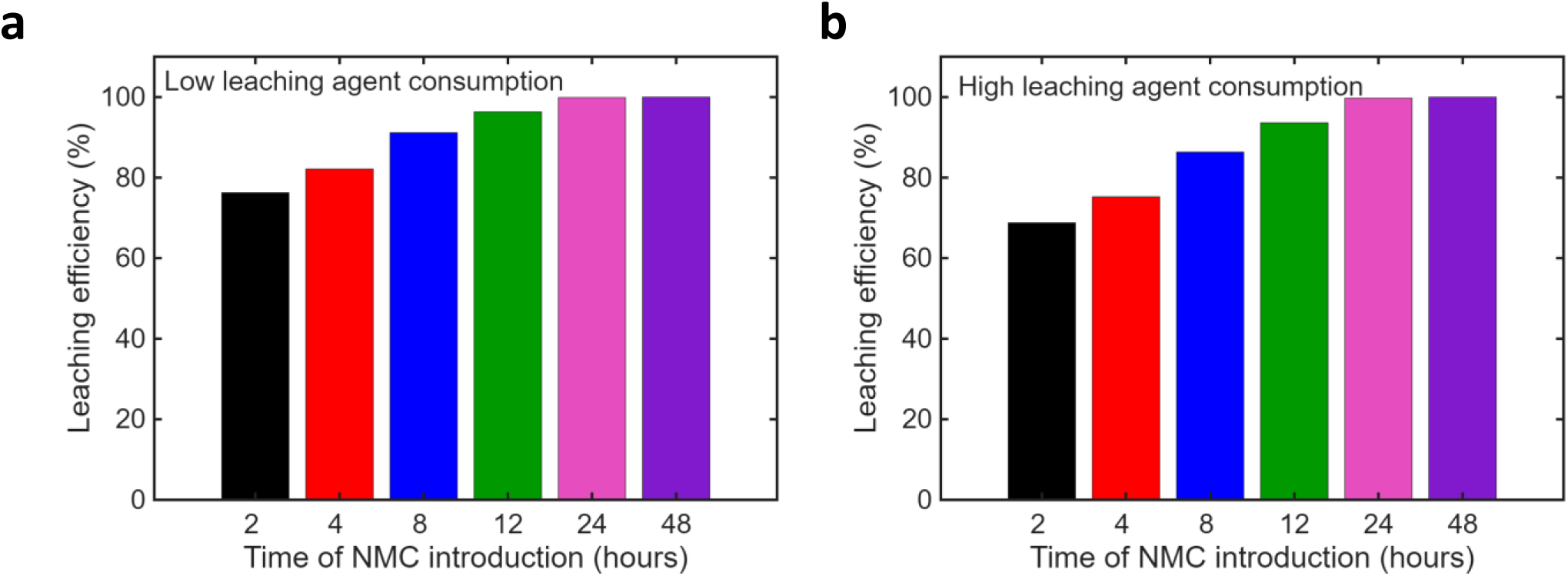
Simulated leaching efficiency after 48 hours of leaching is shown for cultures initiated with the same cell density but with different times of NMC introduction. All the parameters match the values listed in Table 3, except the consumption rate of the leaching agent when leaching NMC. (a) Low leaching agent consumption rate, *s* = 10^3^ U/mg. (b) Low leaching agent consumption rate, *s* = 10^5^ U/mg.

## Notes

### Competing Interest Statement

The authors have declared no competing interest.

